# ARCADE: Controllable Codon Design from Foundation Models via Activation Engineering

**DOI:** 10.1101/2025.08.19.668819

**Authors:** Jiayi Li, Hong-Sheng Lai, Litian Liang, Shiyi Du, Shijie Tang, Carl Kingsford

## Abstract

Codon sequence design is crucial for generating mRNA sequences with desired functional properties for tasks such as developing mRNA vaccines or gene editing therapies. Yet existing methods lack flexibility and controllability to adapt to various design objectives. We propose a novel machine learning-based framework, ARCADE, that enables flexible and controllable multi-objective codon design. Leveraging inherent knowledge from pretrained genomic language models, ARCADE extends activation engineering, a technique originally developed for controllable text generation, beyond discrete feature manipulation such as concepts and styles, to steering continuous-valued biological metrics. Specifically, we derive biologically meaningful semantic steering vectors in the model’s activation space, which directly control properties such as the Codon Adaptation Index, Minimum Free Energy, and GC content. Experimental results demonstrate the flexibility of ARCADE in designing codon sequences with multiple objectives, underscoring its potential for advancing programmable biological sequence design. Our implementation is available at https://github.com/Kingsford-Group/arcade.

## 1 Introduction

Codon design aims to construct mRNA sequences that encode a given protein while enabling customized control over multiple key properties, such as codon usage bias, secondary structure stability, and desired protein abundance levels. These properties collectively influence mRNA translation efficiency, stability, immunogenicity, and have direct implications in developing mRNA-based vaccines and protein therapeutics [28,14,47,34].

Most existing codon design methods focus on a limited set of objectives such as codon usage bias (Codon Adaptiveness Index, CAI) or RNA structural stability (Minimum Free Energy, MFE). Few offer flexible selection and control of multiple sequence properties. Machine learning (ML)-based methods [15,40] exhibit the ability to accommodate diverse design goals without hand-crafted design criteria over rule-based algorithms [16,49]. However, this flexibility reflects training-data skewness toward objectives such as high CAI or low MFE, without control over weights of different objectives. They also often require computation-intensive retraining to target new design goals.

Originally proposed for controllable text generation in language models (LMs), activation engineering uses the LM’s inherent knowledge, modifying the LM’s internal activations during inference to steer model outputs towards desired properties [44,46]. Without objective-specific retraining for varying design requirements [7,10,21,25,38], activation engineering promises to be a solution to controllable codon design.

Still, there are two major hurdles in the development of controllable codon design methods. First, in natural language processing, controllable generation typically steers discrete and human-interpretable properties, such as sentiment, topic, or styles [21,30,25,10,26]. However, properties in genomics, such as biological functionality (e.g., protein expression), structural stability, and translation rate, are often continuous and require computational models or experimental assays for quantification. This highlights a need to extend controllable generation methods to control over continuous properties for genomics sequence design tasks.

Second, current genomic language models (gLMs) [28,23,9,19,31] largely adopt encoder-only architectures, which learn rich representations useful for various predictive tasks but do not output raw sequences as decoder-only LMs do. This necessitates additions to gLMs to facilitate the integration of controllable generation methods.

We propose ARCADE (Activation engineeRing for ControllAble coDon dEsign), an ML-based framework that enables fine-grained and multi-objective control over codon design. ARCADE is designed as a controllable design framework rather than an optimization methodology, enabling guided modulation of and adaptation to various biological properties such as CAI, MFE, GC content, and their combinations. ARCADE leverages activation engineering on pretrained gLMs by constructing and injecting biologically meaningful semantic steering vectors into a pretrained gLM’s latent activation space, allowing direct and adaptable control over diverse continuous attributes. This capability opens the door to rapid exploration of design objectives, scalable generation of biologically coherent variants, and multi-objective codon design tasks using a unified and efficient framework.

Our contributions are as follows:

i. We developed a novel ML-based framework for controllable codon sequence design, enabling flexible steering across multiple sequence properties without updating model weights.
ii. We enabled encoder-only gLMs to perform controllable codon design by introducing a decoding proxy that maps representations to codon sequences.
iii. We established a generalizable activation engineering framework capable of handling continuous-valued biological properties, creating biologically meaningful steering vectors that enable explicit control over key biological attributes such as protein expression, mRNA structure stability, and codon usage.
iv. We extensively validated our method on various codon design objectives and their combinations. It fills the gap in existing ML frameworks to support multi-objective codon design with fine-grained and explicit control over each objective.

## 2 Related work

### Controllable text generation

Controllable text generation enables the steering of language model outputs in terms of style, content, or factuality. CTRL [25] introduces control codes in conditional transformers. Li et al. [29] propose prefix tuning, which enables lightweight adaptation by optimizing small continuous prompts. PPLM [10] offers inference-time control using external attribute classifiers.

Subramani et al. [44] and Hernandez et al. [17] develop activation-based methods that modify internal representations to reflect specific knowledge or behavior.

Meanwhile, activation engineering techniques have gained popularity for their efficiency and flexibility by modulating internal activations of pretrained models without gradient-based optimization or parameter updates. ActAdd [46] steers generation using the activation difference between prompt pairs. CAA [41] refines this using the average of the activation differences between contrastive prompt sets to control hallucination and text sentiment. Konen et al. [26] allow nuanced control of text style by averaging activations from examples with desired stylistic attributes. These approaches mark significant progress toward efficient, flexible, and precise text generation control, though none have been applied to the codon design task and biological sequence generation in general.

### Genomic language models

Advancements in large language models (LLMs) have spurred the development of gLMs, which are large-scale pretrained foundation models capable of performing multiple tasks in understanding and interpreting biological sequences, particularly DNA and RNA sequences. For example, DNABERT [23], Nucleotide Transformer [9], HyenaDNA [36], and Evo [35,6] are all large-scale gLMs for DNA sequence modeling.

For RNA sequence modeling, gLMs like RNABERT [1] and BigRNA [8] are trained on general RNA sequences, while Li et al. [28] developed a gLM that is more suitable for mRNA and protein expression-related tasks by using the codons of RNA sequences as input. We leverage the representation power of gLMs to extract embeddings of codon sequences.

### Codon design

Early computational methods for codon design focus on matching the codon usage frequency of the target gene to that of the host organism. For instance, JCAT [16] greedily selects the most frequent synonymous codons in the identified highly expressed genes of the host. More recent algorithms address the complexity and multi-objective nature of codon design. LinearDesign [49] uses lattice parsing to navigate the search space for possible codon sequences to jointly optimize the CAI and MFE, but its decoding complexity scales quadratically with the input length, making it computationally expensive to design long sequences. These non-machine learning methods lack flexibility and generalizability, and can be time-consuming.

Machine learning approaches learn latent features directly from sequences to guide codon design, offering efficient inference supported by GPU acceleration. Fu et al. [15] used a BiLSTM-CRF network and treated codon design as a sequence annotation problem, where they mapped amino acid sequences to codon sequences based on the codon usage patterns in the host genome. CodonBERT [40], a BERT-based architecture, applies cross-attention between amino acid and codon sequences to learn contextual codon preferences. GEMORNA [48] is a deep generative model that designs complete mRNA sequences, jointly generating coding sequences and untranslated regions with transformer-based architectures. RNADiffusion [22], a latent diffusion model that maps RNA sequences to a low-dimensional latent space, enables guided generation by leveraging gradients from reward models operating on the same latent space.

## 3 Methods

### 3.1 The codon design problem

In this work, the codon design problem is defined as rewriting a given input codon sequence to promote specific biological properties while maximally preserving synonymous fidelity and the biological integrity of the input sequence. Formally, let 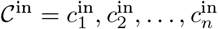 be an input codon sequence, where each codon 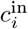 translates to amino acid *ai* via the translation function 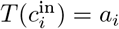. The objective is to construct an output sequence 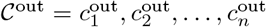, such that 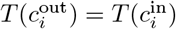 for each *i*, and 𝒞^out^ exhibits more favorable biological properties. This latter goal is a multi-objective function that depends on the use case and users’ desires, which are manifested by the user selecting a set of properties *P* for which they want to enhance. The varied properties in Section 3.7 provide examples of these objective functions.

### 3.2 Framework overview

To solve the codon design problem defined in Section 3.1, we propose to leverage activation engineering to manipulate internal activations within pretrained foundation models and steer the output sequences toward user-specified directions of biological properties.

Specifically, as illustrated in Fig. 1, our framework consists of two stages:

**Fig. 1:**
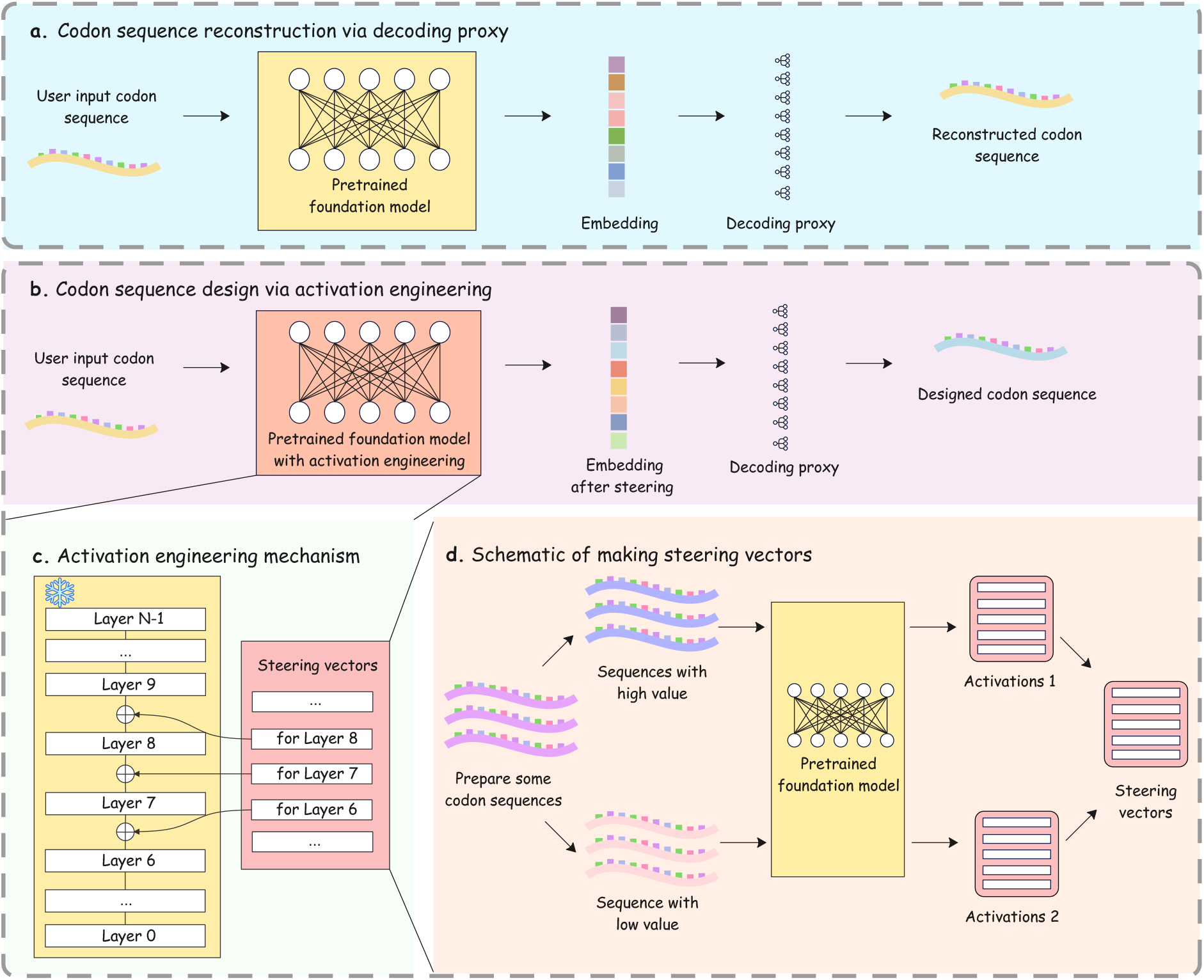
Overview of ARCADE. Given an input codon sequence, **a**. when no steering is applied, the foundation model reconstructs the sequence via a decoding proxy. **b, c**. To steer codon sequences toward desired properties, activation engineering is performed by injecting the constructed steering vectors into the layer activations of a frozen foundation model. **d**. Steering vectors are constructed from the activation differences between high- and low-value examples mutated from a set of seed sequences and scored with a property-specific function.

i. We construct a decoding proxy that equips the encoder-only foundation models with the ability to generate codon sequences through a token-classification head, ensuring faithful reconstruction of codon sequences from their internal representations.
ii. We then derive property-specific steering vectors and perform activation engineering to enable smooth and interpretable control toward user-specified biological objectives.

The following sections elaborate on each component in detail: Section 3.3 describes the activation engineering principle and the construction of steering vectors, and Section 3.4 details the token classification approach as decoding proxy.

### 3.3 Activation engineering for codon design

To enhance a set of user-specified biological properties *P*, we apply activation engineering (Fig. 1**b**). For an input sequence *x*, we modify the activations of the frozen foundation model to steer its output toward a desired direction, such as increasing or decreasing the value of a specified property in P.

Formally, to steer towards properties in *P*, a steering vector 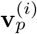 is derived for each layer *i* and each property *p* ∈ *P*. A linear combination of these vectors is added to the activations **a**^(*i*)^(*x*) at layer *i* (Fig. 1**c**):

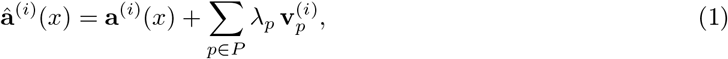

where **a**^(*i*)^(*x*) denotes the original activation at layer *i*, 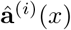 is the modified activation, and the *λp* values control the steering strengths for each property.

To construct the steering vector 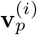 for a property *p* (Fig. 1**d**), we label sequences with high values of the property *p* by 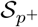 and low values as 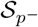. We denote by 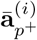 and 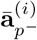 the mean activation vectors at layer *i* for sequences in 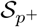 and 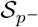, respectively. The steering vector toward a higher value of the property *p* is defined as

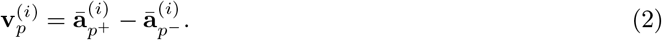

To obtain 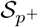 and 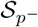 samples for each property *p*, we propose a data-driven construction method tailored for continuous-valued biological attributes. Specifically, starting from a small set of seed codon sequences, we introduce property-specific directed mutations to generate sequence variants with high and low values of that property, forming 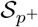 and 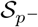, respectively. The mutation method depends on the property of interest (see Appendix A.2 for details). In practice, we obtain the seed set by randomly sampling only 1% of coding sequences from the training set (see Section 3.6), yielding 77 sequences with matched length distribution. Despite its small size, this seed set provides sufficiently robust activation statistics for steering vector computation.

The effectiveness of this approach can be understood through the lens of representation geometry [39]. Pretrained genomic foundation models are expected to encode biologically meaningful embeddings within their latent activation spaces [46]. Different biological properties (e.g., GC content, CAI) correspond to distinct directions or manifolds in this space. By contrasting activations of samples with high and low property values, we can approximate the “direction” along which this property varies. For continuous biological metrics, activation-based modulation captures the linear approximation of the property’s gradient in the latent space. Thus, injecting the corresponding steering vector during inference shifts model activations toward specific directions (i.e., higher or lower values of that property).

As a result, the steering operation can modulate properties through manipulation of internal activations, without modifying model parameters, enabling gradient-free, property-aware, and fine-grained control over diverse biological design objectives.

### 3.4 Token classification as decoding proxy

While activation engineering is typically applied to language models with decoders that can output raw sequences, gLMs are often encoder-only models without such capability [36,5,23,28]. To adapt an encoder-only model for codon design and enable the output of modified sequences, we design a reconstruction mechanism to translate internal representations to discrete codon sequences. This design aligns with prior controllable generation methods in image editing [33,18,24] and text editing domains [10,27,26,41,46], where generators are trained to modify specific attributes while preserving global semantics or structure. Specifically, we introduce a token classification head, a single linear projection from encoder hidden states to codon logits, that functions as a lightweight decoding proxy(Fig. 1**a**). Thus, the alteration of layer activations will be propagated through the token classification head, resulting in a controlled final output of modified codon sequences.

Let 𝒞 = *c*1, *c*2, …, *cn* denote a codon sequence of length *n*, where each *ci* belongs to the vocabulary 𝒱 of the 64 possible codons. The model encodes the input into contextual representations **H** = **h**^1^, **h**^2^, …, **h**^*n*^, where **h**^*i*^ ∈ ℝ^*d*^ denotes the hidden representation at position *i*. A token classification head maps each **h***i* to a categorical distribution over the possible codons:

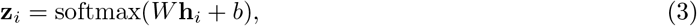

where *W* ∈ ℝ^|𝒱|*×d*^ and *b* ∈ ℝ^|𝒱|^ are learnable parameters. The model is fine-tuned to minimize the cross-entropy loss between the predicted distribution and the ground-truth codon label *yi*:

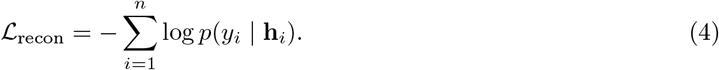

To ensure synonymy, we apply a codon-level mask that restricts predictions to codons encoding the same amino acid as the original input (Appendix A.3). This formulation casts codon reconstruction as a constrained token-level classification task, enabling controllable generation of synonymous codon sequences through subsequent activation steering.

Appending the token classification head on top of the encoder-only foundation model backbone [28], we fine-tune the resulting model using Low-Rank Adaptation (LoRA) [20] (see GENCODE dataset in Section 3.6 and training details in Appendix A.1). The resulting model achieves 99.9% accuracy at the token level in reconstructing input codon sequences. We freeze all the model parameters, and use this fixed model for subsequent sequence design experiments. Notably, this decoding proxy is only required for encoder-only backbones; when ARCADE is applied to generative gLMs with native decoding capability, no proxy is needed. We demonstrate this setting in Section 4.5.

### 3.5 Foundation model backbones

We build ARCADE on two foundation backbones: a codon-level model used for most experiments, due to its natural fit to our design task, and a single nucleotide-level model to demonstrate the generalizability of our framework across backbone architectures.

The codon-level backbone was introduced by Li et al. [28], a BERT [11] encoder pretrained on over 10 million mRNA codon sequences spanning diverse species, including mammals, bacteria, and human viruses. We use its publicly released checkpoint and tokenizer. The model uses a 12-layer transformer encoder with 12 self-attention heads per layer and a hidden size of 768. Input sequences are tokenized at the codon level using a 64-token vocabulary representing the standard genetic codons. The model is pretrained using a masked codon modeling objective to capture contextual patterns of codon usage across diverse organisms.

We use Evo [35], a single nucleotide-level backbone, as an example to assess ARCADE’s generalizability. Evo is a state-of-the-art genomic language model (gLM) with approximately 7 billion parameters. Since its architecture uses a decoder, we remove the token classification head and use its released checkpoint for inference.

### 3.6 Datasets

Our main dataset is constructed by extracting protein-coding transcripts from the GENCODE Human Release 47 (GRCh38.p14) annotation [13] (abbreviated “GENCODE”). The coding regions (CDSs) are isolated from 85,099 transcripts. To conform to the input length limit of 1,024 codons of the foundation model [28], we filter out sequences longer than 3,072 base pairs. The dataset is split into training and testing sets at a 9:1 ratio, while ensuring all transcript variants of the same gene are assigned to the same split to prevent information leakage between transcripts of the same gene.

The training set is used both to fine-tune the token classification head and to compute activation statistics for steering vector construction, while the held-out test set contains 8,508 sequences and is used for evaluation on all related experiments.

For additional evaluation on experimentally measured monomeric Red Fluorescent Protein (mRFP) expression, we use the data from Nieuwkoop et al. [37], which contains 1,459 variants of the mRFP gene annotated with their measured expression levels. This benchmark enables us to assess the effectiveness of steering methods on a functionally grounded biological property.

### 3.7 Diverse codon properties

To evaluate whether ARCADE generalizes across diverse biological and functional properties of codon sequences, we applied it to generate sequences that optimize a wide range of metrics. Details of the metric calculations are provided in Appendix A.8.

#### Codon usage metrics

The *Codon Adaptation Index (CAI)* is widely used to evaluate how closely a codon sequence aligns with the preferred codon usage in a given organism [43]. CAI values range from 0 to 1 (unitless), with higher values implying greater compatibility with the host’s translational machinery. Optimizing CAI has been shown to enhance the efficiency in protein translation [51,2].

#### Structure-related metrics

*Minimum Free Energy (MFE)* [12], measured in kcal/mol, reflects the thermodynamic stability of mRNA folding. We report *Negated Minimum Free Energy (NMFE)* in the results, the additive inverse of MFE. Thus, higher NMFE values correspond to more stable mRNA structures. *GC content* measures the fraction of guanine and cytosine nucleotides in the sequence (range 0–1). Higher GC content is typically correlated with increased structural stability [50].

#### Immunity-related metrics

CpG and UpA dinucleotide densities are known to affect innate immune responses and vaccine safety [42]. The *CpG density* and *UpA density* metrics are computed as the number of CG or UA dinucleotides per 100 base pairs(bp) of sequence [4].

#### Functional outputs

The expression level of mRFP (*mRFP expression*) serves as a proxy for evaluating the functional impact of codon design strategies. Increasing the expression of a gene (such as mRFP) is a typical goal for improving the efficacy of gene therapies [3,37]. The details of calculating mRFP values for the computationally generated sequences are presented in Appendix A.8.

## 4 Experiments

We demonstrate that ARCADE enables effective and flexible control over a range of attributes for codon design (Fig. 2). We evaluate controllability in two settings: single-property steering, which modulates individual biological metrics, and multi-objective steering, which includes both paired and triple objectives to demonstrate ARCADE’s ability to steer multiple properties.

**Fig. 2:**
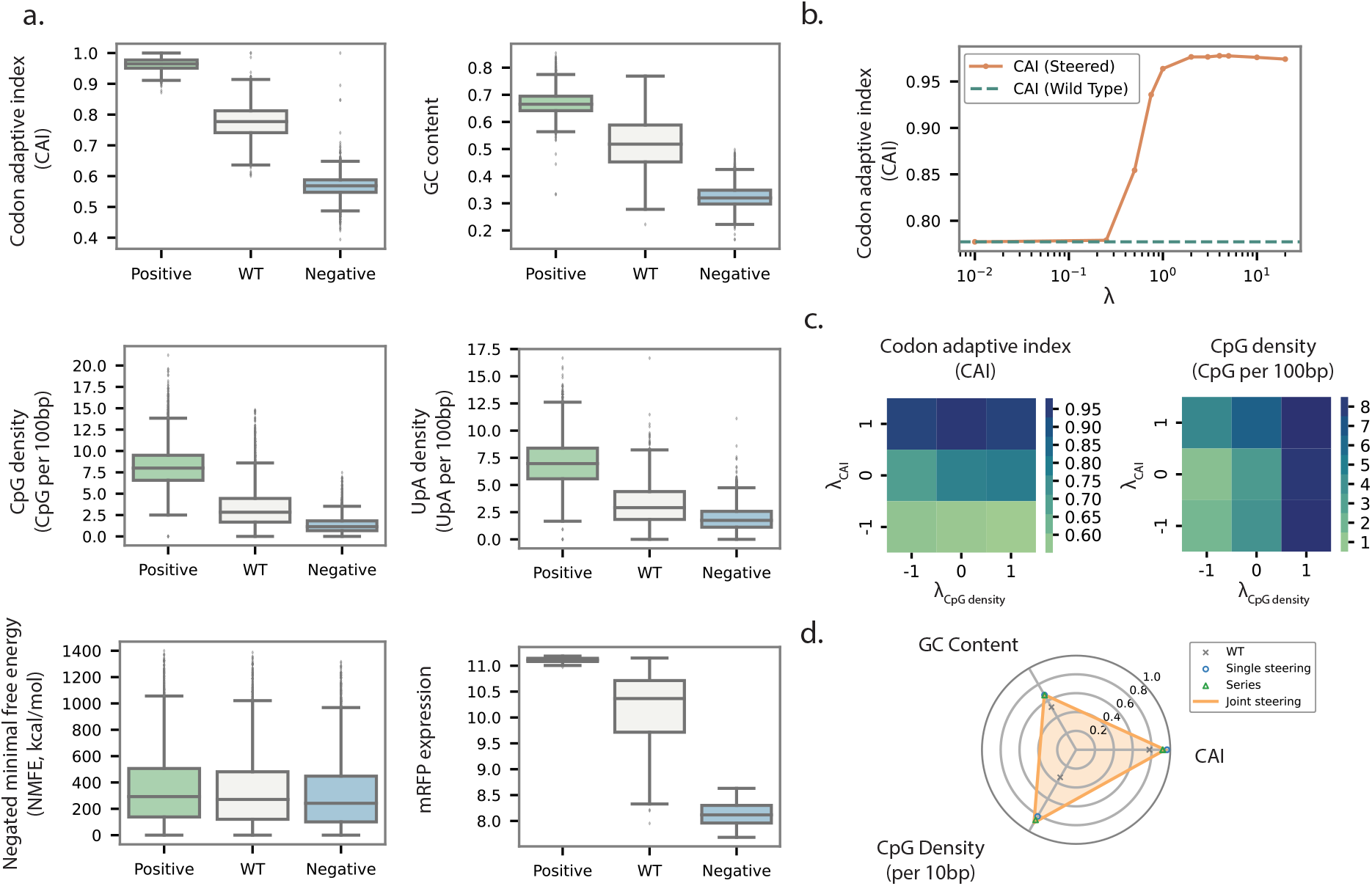
**a**. Single-property steering. Boxplots show distributions of codon properties under positive steering, negative steering, and wild-type (WT) sequences. Using steering strength *λ* = 1, ARCADE consistently shifts metrics (CAI, MFE, GC content, CpG density, UpA density, and mRFP expression) in the expected directions. **b**. Effect of steering strength on CAI. CAI values under steered (blue) and wild-type (red) sequences are plotted against steering strength *λ* (log scale). CAI increases rapidly for *λ* ∈ [0.5, 4] and plateaus for larger *λ*, indicating stable controllability without over-amplification. The exact values for each point are provided in Table 6 in Appendix A.5. **c**. Paired-property steering. Heatmaps show mean values of CAI (left) and CpG density (right) under *λ*CAI, *λ*CpG ∈ {−1, 0, +1}. Positive values increase the corresponding property, negative values decrease it, and zero corresponds to no steering. **d**. Triple-property steering. Radar plot comparing WT, single-property steering, sequential (series) steering, and joint steering across CAI, GC content, and CpG density. In series steering, steering vectors are applied sequentially (first CAI, then GC, then CpG), whereas joint steering combines all three vectors simultaneously using equal weights (*λ* = 1 for each property).

### 4.1 Steering single properties

Adding and subtracting the steering vectors (“positive” and “negative” steering) for one property consistently shifted the distribution of the property in the correct direction across CAI, GC content, CpG density, UpA density, MFE, and mRFP expression (Fig. 2**a**, Table 4 in Appendix; mRFP expression is evaluated by the prediction model, see Appendix A.8 for details). Each inference run on the GENCODE test set (Section 3.6) requires approximately 20 minutes on an NVIDIA A100 GPU with 40 GB of memory. Table 5 (in Appendix) presents the results of the one-sided t-tests comparing steered sequences with the wild-type baseline. This confirms that ARCADE enables robust bidirectional control of individual attributes and has applicability across multiple aspects of codon design objectives.

We next examine the effect of steering strength *λ* (i.e., *λ*_*p*_ defined in Eq. 1) using CAI as an example (Fig. 2**b**). As *λ* increases, we observe a classic monotonic dose-response curve. The regime *λ* ∈ [0.5, 4] is sufficient to meaningfully influence sequence perturbation yet without exceeding the model’s representation capacity, providing practical guidance for choosing *λ*. These results indicate that ARCADE offers fine-grained and stable modulation without over-amplification.

### 4.2 Steering multiple properties

Flexible steering of multiple properties is important for practical applications where codon sequences must satisfy multiple biological constraints, such as increasing expression while maintaining structural stability. Thus, we further examine ARCADE’s capacity for multi-objective codon design by jointly steering multiple sequence properties. We found that jointly steering two different properties (e.g., CAI and CpG density) by linearly combining the respective steering vectors elicited desired changes in both attributes (Fig. 2**c**). We also demonstrate ARCADE’s effectiveness in simultaneously steering towards the property pairs CAI + NMFE, CAI + GC content, and mRFP + NMFE in Table 1, where joint steering improves both target metrics relative to the wild-type (WT) baseline and matches or exceeds results by single-property steering on each individual metric.

**Table 1:** Joint steering across multiple objective pairs. Each row block corresponds to a pair of target properties. “Single” steers toward a single objective (*λ* = 1). “Joint Steering” jointly steers the two directions, both using *λ*_*p*_ = 1 (Eq. 1). For the mRFP + NMFE^*^ objective pair, both metrics are evaluated on the mRFP dataset; NMFE^*^ is evaluated on mRFP sequences.

| Objective Pair | Metric | WT | Single | Joint Steering |
| --- | --- | --- | --- | --- |
| CAI + CpG | CAI | 0.778 | 0.964 | 0.937 |
|  | CpG | 3.376 | 8.168 | 8.479 |
| CAI + NMFE | CAI | 0.778 | 0.964 | 0.978 |
|  | NMFE | 331.09 | 350.42 | 424.05 |
| CAI + GC | CAI | 0.778 | 0.964 | 0.978 |
|  | GC | 0.522 | 0.671 | 0.672 |
| mRFP + NMFE* | mRFP | 10.162 | 11.110 | 11.144 |
|  | NMFE* | 195.97 | 223.37 | 225.93 |

To better understand the effectiveness of multi-objective steering, we analyze the geometry of steering directions in terms of metric values. As shown in Fig. 3, joint steering allows for navigating the trade-off between the two metrics, CAI and NMFE. Instead of collapsing towards one objective, the model responds to joint steering by exploring a shared region of the representation space where both objectives can improve.

**Fig. 3:**
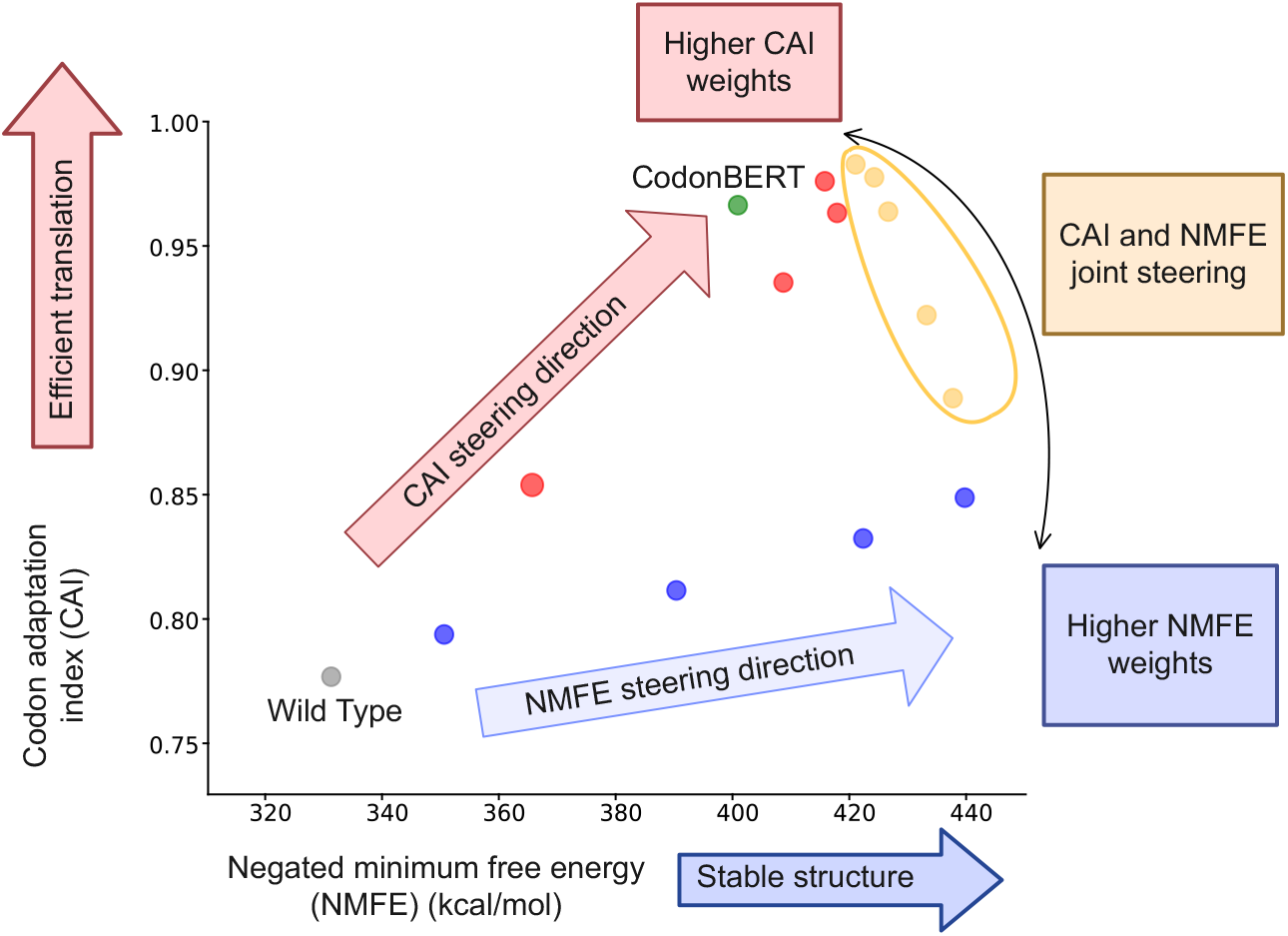
Visualization of the combined steering effect. Points represent average value shifts steered by ARCADE across multiple steering strengths *λ* along CAI (red), NMFE (blue), and their combination (CAI + NMFE, yellow). The resulting geometry suggests that the combination of individual steering creates a controllable region in the model’s latent space, where multi-objective trade-offs can be navigated via vector-based interpolation. See Appendix A.7 for raw values.

Extending the pairwise scheme to three properties, we compare two strategies for multi-objective steering: (i) joint steering, where all three objectives are optimized simultaneously; and (ii) series steering, where the steering vectors for CAI, GC content, and CpG density are applied sequentially. As shown in Fig. 2**d**, both strategies yield comparable improvements across all three metrics relative to single-property steering. Joint steering is generally preferred since the effectiveness of earlier-applied objectives in the series setting slightly diminishes due to accumulated perturbations, though the degradation remains minor (Table 8 in Appendix). Together, these results establish ARCADE as a general framework for flexible and robust multi-objective codon design.

### 4.3 Ablation studies

To verify that the observed effects are attributable to meaningful steering directions rather than mere perturbation, we compare steering on CAI and NMFE with random steering vectors (“Random”) and with steering strength *λ* = 0. The random steering vector is constructed to have the same shape and magnitude as the derived steering vector (*λ* = 1, Section 3.3) but filled with random values. The two ablations result in values nearly identical to those of the wild-type sequences, within 0.01% relative change (Table 2). These results confirm that the observed effects do not arise from arbitrary perturbations or noise, corroborating that our steering directions are both informative and closely linked to the biological properties of interest.

**Table 2:** Ablation study. “Random” applies random steering vectors with the same *ℓ*_2_ norm of the corresponding *λ* = 1 steering vectors. “*λ* = 0” applies no steering at all. “*λ* = 1” applies the steering vectors for each property with steering strength set to 1.

|  | CAI | NMFE | GC content |
| --- | --- | --- | --- |
| WT | 0.778 | 331.09 | 0.522 |
| Random | 0.778 | 331.12 | 0.522 |
| $\lambda = 0$ | 0.778 | 331.12 | 0.522 |
| $\lambda = 1$ | 0.964 | 350.42 | 0.671 |

### 4.4 ARCADE as a unified approach to codon design objectives

ARCADE serves as a unified framework capable of steering diverse codon design objectives, and we evaluate its effectiveness by comparing ARCADE against several widely used and state-of-the-art codon design methods described in Section 2. Most existing methods support a limited set of objectives. For example, JCAT [16] only optimizes for CAI; CodonBERT [40] and BiLSTM-CRF [15] only target CAI and MFE, yet they do not allow control of the trade-off between the two metrics. RNADiffusion [22] is not included in our comparisons due to no official implementation or publicly available pretrained model exists.

Though ARCADE is designed for flexible control, it is competitive with these baseline methods on the metrics for which they aim to optimize (Tables 3**a** and 3**b**). The joint steering model CAI+NMFE steers both properties simultaneously with *λ* = 1 and achieves competitive or superior results on both metrics compared to the other methods.

**Table 3:** Comparison of ARCADE with other methods. “unit NMFE” is the NMFE normalized by sequence length. The results of the best method are marked in bold. **a**. is the comparison with BiLSTM-CRF **b**. is performed on the subset of sequences with length *<* 1500 to accommodate its input length constraint. When comparing with LinearDesign **c**. on mRFP expression, ARCADE applies steering only on this metric. In comparison with GEMORNA **d**., ARCADE jointly steers CAI, NMFE, and GC content, all with strength *λ* = 1. In all other comparisons, ARCADE jointly steers CAI and NMFE, both with *λ* = 1.

| a. Comparison with CodonBERT and JCAT |  |  |  |  | b. Comparison with BiLSTM-CRF |  |  |  |
| --- | --- | --- | --- | --- | --- | --- | --- | --- |
|  | WT | ARCADE | CodonBERT | JCAT |  | WT | ARCADE | BiLSTM-CRF |
| CAI | 0.773 | 0.978 | 0.967 | **1.0** | CAI | 0.778 | **0.979** | 0.750 |
| NMFE | 331.09 | **424.05** | 400.71 | 369.18 | NMFE | 220.26 | **362.20** | 245.56 |
| unit NMFE | 0.324 | **0.413** | 0.399 | 0.363 | unit NMFE | 0.322 | **0.411** | 0.362 |

| c. Comparison with LinearDesign |  |  |  | d. Comparison with GEMORNA |  |  |
| --- | --- | --- | --- | --- | --- | --- |
|  | ARCADE | CodonBERT | LinearDesign |  | ARCADE | GEMORNA |
| mRFP expression | **11.186** | 9.916 | 9.570 | CAI | **0.969** | 0.878 |
| CAI | **0.977** | 0.964 | 0.944 | NMFE | **421.50** | 384.33 |
| NMFE | 416.30 | 395.46 | **568.172** | GC Content | **0.672** | 0.608 |

**Table 4:** ARCADE effectively steers individual metrics as described in Section 3.7. “WT” denotes the unmodified wild-type input sequences. mRFP expression is evaluated on the dataset from Nieuwkoop et al. [37], and all other metrics are averages over GENCODE test set (see Section 3.6). (Steering strength *λ* = 1 for all experiments.)

|  | Translation (CAI) | Stability (NMFE) | GC content | CpG density | UpA density | mRFP expression |
| --- | --- | --- | --- | --- | --- | --- |
| Positive | 0.964 | 350.42 | 0.671 | 8.168 | 7.024 | 11.110 |
| WT | 0.778 | 331.09 | 0.522 | 3.376 | 3.159 | 10.162 |
| Negative | 0.568 | 305.22 | 0.325 | 1.330 | 1.915 | 8.129 |

**Table 5:** One-sided t-test results for single-metric steering. We report the t-statistics (*t*) and their corresponding p-values (*p*). Each test compares the metric values of all steered sequences in the test set (positive or negative) against their wild-type (WT) counterparts. The t-statistic (*t*) indicates the direction and magnitude of the difference relative to wild-type, with positive values showing higher values in positively steered sequences and negative values showing lower values in negatively steered sequences. The p-value (*p*) represents the probability of observing such a difference under the null hypothesis, with smaller values indicating higher statistical significance.

|  | CAI |  | GC content |  | CpG density |  | UpA density |  | NMFE |  | mRFP expression |  |
| --- | --- | --- | --- | --- | --- | --- | --- | --- | --- | --- | --- | --- |
| | $t$ | $p$ | $t$ | $p$ | $t$ | $p$ | $t$ | $p$ | $t$ | $p$ | $t$ | $p$ |
| Positive | 336.81 | 0 | 145.11 | 0 | 135.65 | 0 | 135.66 | 0 | 4.84 | $6.62 \times 10^{-7}$ | 19.76 | $6.73 \times 10^{-51}$ |
| Negative | -334.60 | 0 | -193.09 | 0 | -75.36 | 0 | -58.41 | 0 | -6.63 | $1.77 \times 10^{-11}$ | -40.84 | $1.20 \times 10^{-113}$ |

**Table 6:** Performance of CAI steering with different steering strength *λ*.

| Metric | WT | $\lambda = 0$ | $\lambda = 0.25$ | $\lambda = 0.5$ | $\lambda = 0.75$ | $\lambda = 1$ | $\lambda = 2$ | $\lambda = 3$ | $\lambda = 4$ | $\lambda = 5$ | $\lambda = 10$ | $\lambda = 20$ | CodonBERT |
| --- | --- | --- | --- | --- | --- | --- | --- | --- | --- | --- | --- | --- | --- |
| NMFE | 331.09 | 331.12 | 331.29 | 365.49 | 408.50 | 417.69 | 415.59 | 405.43 | 401.27 | 399.46 | 396.58 | 394.47 | 400.71 |
| CAI | 0.7770 | 0.7780 | 0.7790 | 0.8544 | 0.9358 | 0.9638 | 0.9765 | 0.9765 | 0.9777 | 0.9776 | 0.9760 | 0.9740 | 0.9669 |

We also compared ARCADE with LinearDesign [49], a recent non-machine learning method for jointly optimizing CAI and MFE. For fairness, we adopt their combination mode (with their parameter *λ* = 4, distinct from our method). Due to the high computational cost of LinearDesign, we evaluate the methods in Table 3**c** on a randomly selected subset of 100 coding sequences from the full test set (size 8, 508).

As shown in Table 3**c**, all methods achieve comparable CAI values, while LinearDesign produces the most stable sequences in terms of NMFE. However, LinearDesign is inherently limited to optimizing structural metrics such as CAI and NMFE and cannot directly account for protein-specific expression levels. In contrast, ARCADE flexibly accommodates diverse objectives by steering sequence representations toward user-specified properties. When evaluated on mRFP expression, ARCADE attains higher predicted expression than both LinearDesign and CodonBERT. At the same time, ARCADE maintains competitive performance on CAI and NMFE when jointly steering this pair of objectives, comparable to methods specialized for optimizing these properties. Although ARCADE is not an optimization algorithm, it enables flexible exploration of the design space while supporting multi-objective steering in a unified framework. This result underscores ARCADE’s capacity for flexible control over sequence design objectives, extending beyond CAI and MFE to experimentally measured functional outcomes.

Even for 100 sequences, LinearDesign requires over 8 hours using two Intel(R) Xeon(R) CPU E5-2690v2 3.00GHz (20 cores), highlighting its limitations for large-scale codon design tasks. In contrast, ARCADE enjoys superior scalability, editing these 100 sequences in 16 seconds, including model loading time, using a single NVIDIA A100 GPU with 40GB of memory. Even for 100 sequences, LinearDesign requires over 8 hours (*>* 4.8 minutes per sequence) on two Intel(R) Xeon(R) CPU E5-2690v2 3.00GHz (20 cores), highlighting its limitations for large-scale codon design tasks. In contrast, ARCADE enjoys superior scalability, editing these 100 sequences in 16 seconds (0.16 seconds per sequence) on a single NVIDIA A100 GPU with 40GB of memory.

We further compare ARCADE with GEMORNA [48], a recent machine learning-based method for codon design that generates mRNA sequences directly from protein inputs without offering control over codon-level properties (Table 3**d**). For comparison, we apply ARCADE with joint steering across CAI, NMFE, and GC content, the three metrics reported in the GEMORNA paper. ARCADE attains higher CAI, NMFE, and GC content relative to GEMORNA. These results highlight ARCADE’s ability to simultaneously enhance multiple design objectives, underscoring its advantage as a flexible and general framework for multi-objective codon design.

### 4.5 Extending ARCADE beyond codon-specific foundation models

To assess the generalizability of ARCADE, we explore its application potential on genomic language models (gLMs) beyond the codon-specific architecture [28] that we used in previous experiments (see methods in Section 3.5). We apply ARCADE to Evo [35], a large-scale gLM that learns multimodal representations across genomes. Applying single-property steering on Evo (Fig. 4) shows that ARCADE consistently elicits controllability in the attribute of interest. Each inference run on the GENCODE test set (Section 3.6) this larger model requires approximately 500 minutes on an NVIDIA A100 GPU with 40 GB of memory. These results suggest ARCADE’s potential applicability to a broader range of genomic sequence design tasks, including regulatory elements, non-coding RNA, and protein design.

**Fig. 4:**
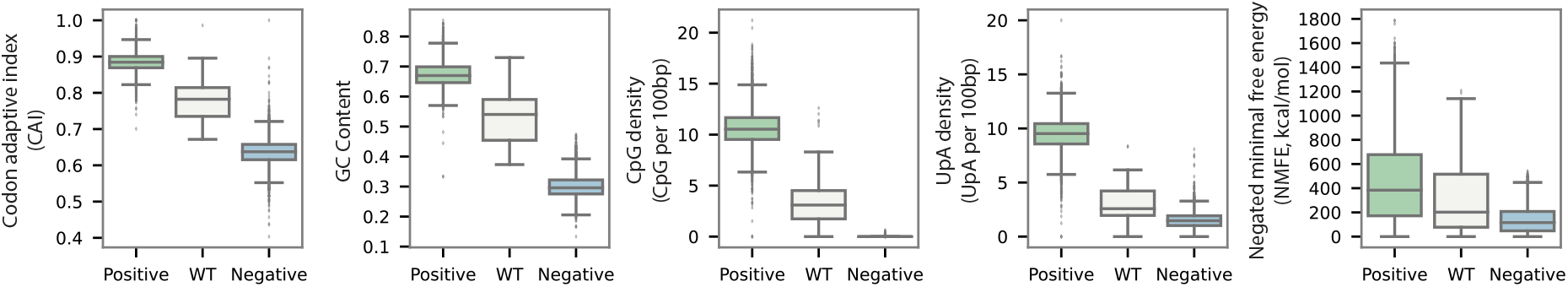
ARCADE applied to Evo demonstrates consistent controllability. Boxplots show distributions of codon properties under positive steering, negative steering, and WT sequences. Steering consistently shifts metrics (CAI, MFE, GC content, CpG density, and UpA density) in the expected directions. Metrics are reported as averages over the GENCODE test set (see Section 3.6). (*λ* = 1.)

## 5 Discussion

We introduce ARCADE, a novel machine learning-based framework for controllable codon design. ARCADE provides a flexible framework for codon design by leveraging activation engineering on pretrained biological foundation models to steer and control multiple sequence-level properties without retraining. It delivers flexible and fine-grained control over attributes that affect translation efficiency, stability, and immunogenicity. This capability allows the design of sequence attributes relevant to protein expression, stability, and immunogenicity.

One insight is that the latent space of pretrained codon language models captures biologically meaningful directions. ARCADE exploits this structure to enable flexible control, supporting both single- and multi-objective generation. Even for properties often considered in trade-off (e.g., CAI vs. MFE), ARCADE identifies directions that jointly improve both metrics towards a Pareto-like trade-off space.

Despite its flexibility, ARCADE has some constraints. Its effectiveness depends on the inductive biases of the underlying foundation model and the sequences from which the steering vectors were constructed. As a result, steering is confined to biologically plausible regions of the sequence space, which may be good for some applications but limiting for others. Moreover, ARCADE applies only global, whole-sequence control and does not provide sequence-specific or position-aware editing.

Future work could address these gaps by integrating local feedback mechanisms or gradient-based refinement to support fine-grained, sample-specific control. Integrating ARCADE into iterative design loops with experimental validation would further validate its utility for synthetic biology.

## Acknowledgment

This work was supported in part by the US National Science Foundation [III-2232121] and the US National Institutes of Health [R01HG012470]. We thank Dr. Guillaume Marçais for insightful feedback and valuable suggestions. Conflict of Interest: C.K. is a co-founder of Ellumigen, Inc.

## A Appendix

### A.1 Implementation details of decoding proxy

To enable codon sequence reconstruction using an encoder-only genomic language model, we attach a token-classification head to pretrained backbone released by Li. et al [28]. This head serves as a lightweight decoding proxy, mapping encoder hidden states to codon-level prediction.

#### Architecture of the decoding proxy

The token-classification head is implemented as a position-wise linear classifier. For each codon position *t*, the encoder hidden state **h**_*t*_ ∈ ℝ^*d*^ is projected to a distribution over the codon vocabulary *V* via:

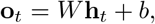

where **o**_*t*_ ∈ ℝ^|𝒱|^ represents the unnormalized logits over the codon vocabulary 𝒱, and *W* ∈ ℝ^|𝒱|*×d*^ and *b* ∈ ℝ^|𝒱|^ are learnable parameters. The logits are converted into a probability distribution

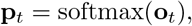

where **p**_*t*_ ∈ ℝ^|𝒱|^ gives the predicted probability of each codon at position *t*. No autoregressive decoding or additional sequence-level modeling is introduced; the proxy performs independent codon-level classification directly from encoder representations. This simple structure allows encoder-level steering to produce predictable changes in the output codons.

#### Fine-tuning setup

To learn this decoding proxy, we fine-tune the model on a codon reconstruction task. Coding sequences from GENCODE v47 (Section 3.6) are truncated or padded to a maximum length of 1,024 codons. Following the setup of Li et al. [28], we keep all backbone parameters frozen and update only (i) the LoRA adapters and (ii) the linear classification head. We use Low-Rank Adaptation (LoRA) with rank 32, *α* = 32, and dropout 0.1. Training is performed with the AdamW optimizer, a learning rate of 5 *×* 10^−5^, batch size 64, and 1,000 optimization steps.

#### Evaluation

For evaluation, we randomly split the full training set described in Section 3.6 into a 98:1:1 train/validation/test partition. We report token-level reconstruction accuracy on the held-out 1% test split. The resulting model achieves 99.9% token-level accuracy in reconstructing input codon sequences, and this fixed model is used for all subsequent sequence design experiments.

### A.2 Deriving 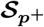 and 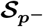 samples via sequence mutations

To construct steering vectors for each metric, we adopt a metric-specific strategy. We randomly sample 1% of coding sequences from the training set, comprising 77 sequences with the same distribution of sequence lengths, and apply controlled mutations to generate sequence variants 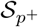 and 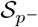.

For CAI, we create 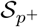 (high CAI) by mutating codons with high usage frequencies, and 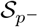 (low CAI) by replacing codons with less frequently used synonyms. A similar strategy is applied for GC content, CpG density, and UpA density.

For NMFE, which is a structural property that is not directly interpretable in terms of codon substitutions, we use LinearDesign [49] to generate high-stability sequences for 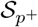 (high NMFE), while 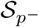 (low NMFE) is generated via random synonymous perturbations.

For the mRFP dataset, which contains mutated coding sequences of the mRFP gene along with experimentally measured expression values, we directly use the top and bottom 3% of sequences ranked by expression as the 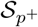 (high-expression) and 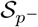 (low-expression) sets, respectively.

### A.3 Synonymous codon mask

To ensure that only synonymous codon variations are considered during reconstruction, we define a binary mask of shape (batch_size, sequence_length, 64) on the output logits of the token classification head by (1) translating the input codon sequence into the corresponding amino acids and (2) marking “1”s at all synonymous codons for each amino acid. The mask is then multiplied to the logits to ensure nonzero output probabilities only for synonymous codons.

### A.4 ARCADE performance across single-property steering tasks

Table 4 reports the results of single-property steering experiments described in Section 4.1. Table 5 reports the results of one-sided t-tests assessing the statistical significance of the steering effects summarized in Table 4. Each test compares the metric distributions of steered sequences (positive or negative) with their wild-type (WT) counterparts across the GENCODE test set.

### A.5 Effect of steering strength *λ*

Tables 6 and 7 show CAI (Fig. 2**c**) and MFE values, respectively, after steering the test set sequences with different *λ*.

**Table 7:** Performance of NMFE steering with different steering strength *λ*.

| Metric | WT | $\lambda = 0$ | $\lambda = 0.5$ | $\lambda = 1$ | $\lambda = 1.5$ | $\lambda = 2$ | $\lambda = 4$ | $\lambda = 10$ | $\lambda = 20$ | CodonBERT |
| --- | --- | --- | --- | --- | --- | --- | --- | --- | --- | --- |
| NMFE | 331.09 | 331.12 | 331.42 | 350.42 | 390.18 | 422.16 | 438.94 | 439.55 | 440.19 | 400.71 |
| CAI | 0.777 | 0.778 | 0.779 | 0.794 | 0.812 | 0.833 | 0.860 | 0.849 | 0.849 | 0.967 |

**Table 8:**
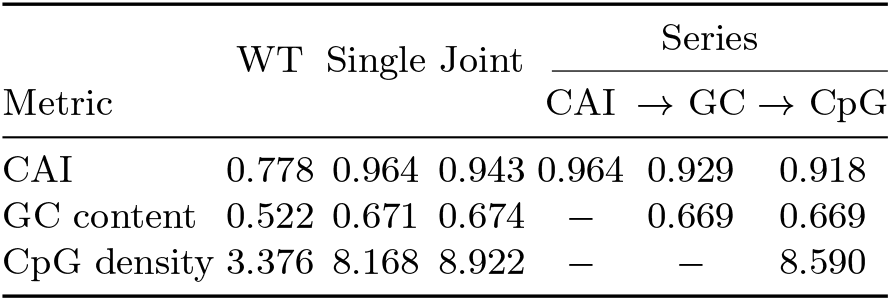
Raw values for triple-property steering. Each column reports mean metric values across the GENCODE test set under different steering configurations. “WT” denotes unmodified wild-type sequences. “Single” corresponds to single-property steering with *λ* = 1 for each property. “Joint” indicates simultaneous steering across CAI, GC content, and CpG density using equal steering strength (*λ* = 1). “Series” denotes sequential steering, where steering vectors are applied in order (CAI → GC → CpG).

### A.6 ARCADE performance across triple-property steering tasks

Table 8 reports the quantitative results underlying the radar plot in Fig. 2**d**.

### A.7 Details regarding Fig. 3

We report the steering strengths represented by the dots in Fig. 3.

For the red dots (CAI steering), from bottom-left to top-right, the steering strengths *λ* are 0.5, 0.75, 1, 2, respectively. The exact value can be referred from Table 6.

For the red dots (NMFE steering), from bottom-left to top-right, the steering strengths *λ* are 1, 1.5, 2, 10, respectively. The exact values are provided in Table 7.

The combination of the *λ*s for the yellow dots, as well as their performance in terms of CAI and NMFE, is presented in Table 9.

**Table 9:** The performance of joint steering in Fig. 3.

| $(\lambda_{\text{CAI}}, \lambda_{\text{MFE}})$ | (1,4) | (1,2) | (1,1) | (4,1) | (10,1) |
| --- | --- | --- | --- | --- | --- |
| MFE | 433.03 | 426.45 | 424.05 | 405.11 | 397.80 |
| CAI | 0.923 | 0.964 | 0.978 | 0.981 | 0.977 |

### A.8 Calculation of metrics

#### Codon Adaptation Index (CAI)

The *CAI* quantifies synonymous codon usage bias across species [43]. It is commonly used as a proxy for translational efficiency. Given an mRNA coding sequence *r* of length |*r*| nucleotides, the number of codons is *n* = |*r*| */*3. Let codon(*r, i*) denoe the *i*-th codon in the sequence, for 0 ≤ *i < n*. Each codon *c* is assigned a relative adaptiveness weight *w*(*c*) ∈ [0, 1], which reflects its frequency relative to the most used synonymous codon for the same amino acid, based on a reference table (e.g., from the Codon Statistics Database [45]).

Then, the CAI of a sequence *r* is defined as the geometric mean of the adaptiveness values over all codons:

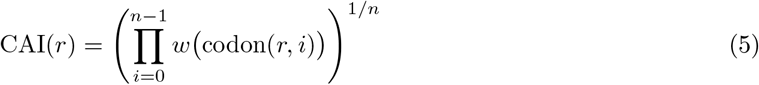

which ranges from 0 to 1, with higher values indicating better alignment with the host’s codon usage preferences [43].

#### Minimum Free Energy (MFE)

The *MFE* of an RNA sequence reflects its predicted thermodynamic stability. A lower MFE indicates a more stable RNA secondary structure. We compute MFE for each mRNA *r* using the `RNAfold` algorithm from the ViennaRNA package (v2.7.0) with default settings [32]. Results are reported in kcal/mol.

To facilitate the presentation, we use the *Negated minimum free energy (NMFE)* as defined in Section 3.7, such that higher NMFE values correspond to greater structural stability.

#### GC content

The *GC content* of an mRNA sequence *r* is defined as the percentage of guanine (G) and cytosine (C) nucleotides it contains. Let #*X*(*r*) denote the count of nucleotide *X* in *r*.

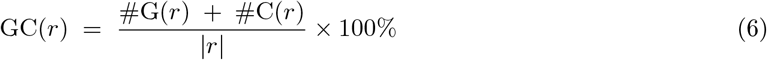

#### CpG density and UpA density

The CpG density is defined as the number of CG dinucleotides per 100 base pairs(bp) [4]. It is calculated as follows:

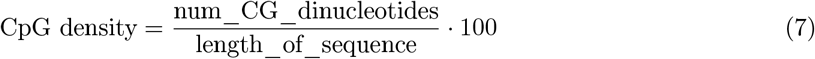

The UpA density is calculated analogously.

#### mRFP expression prediction

Since mRFP expression is an experimentally measured value without a known analytical formula, we train a predictive model by fine-tuning the pretrained model from Li et al. [28]. We attach a regression head (a single linear layer mapping the final hidden representation of each codon to a scalar) on top of the encoder, following the standard HuggingFace token regression setup. The entire model is fine-tuned end-to-end using AdamW with learning rate 5 × 10^−5^, batch size 128, and 200 training epochs, adopting the hyperparameters used in the original work.

Training is performed on the dataset described in Section 3.6 with the original train/validation/test split. The distribution of mRFP expression values in the training set is shown in Fig. 5. We evaluate the model by computing Spearman’s rank correlation between the predicted and experimentally measured expression values, achieving a correlation of 0.87 on the held-out test set.

**Fig. 5:**
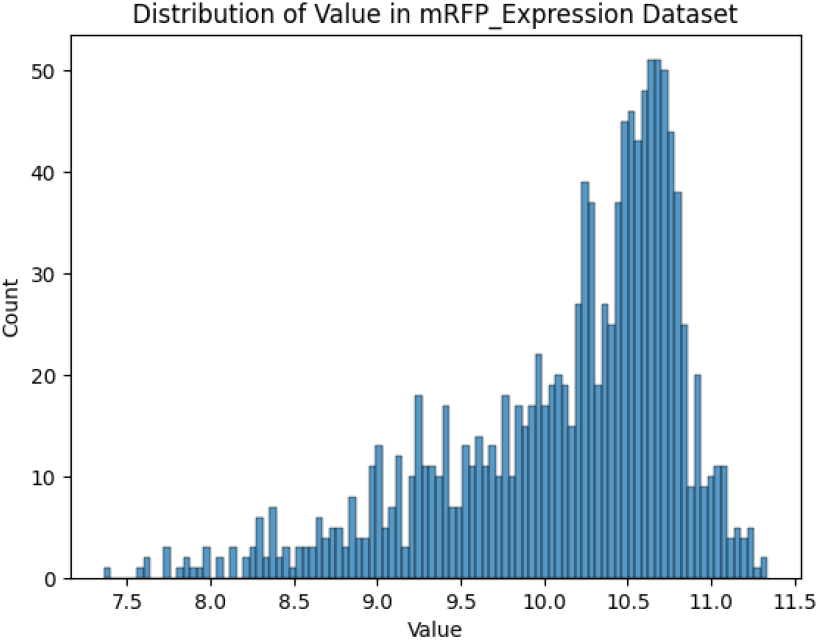
Distribution of mRFP expression.

### B Safety

Our method, ARCADE, introduces controllable codon design by enabling fine-grained manipulation of biological properties such as expression level and structural stability. This can support applications in synthetic biology, gene therapy, and mRNA vaccine development, offering potential societal benefits by accelerating therapeutic design and improving biological sequence performance.

However, the same controllability could raise risks if used to design sequences with unintended or malicious properties, such as avoiding immunogenicity detection or bypassing biosafety checks. Furthermore, because our approach builds on pretrained models, its behavior is ultimately constrained by the biases and limitations of the underlying training data, which may lead to unreliable results in poorly represented sequence regimes.

While this work remains foundational and is not intended for direct deployment, future applications should consider integrating biosafety filters, rigorous interpretability tools, and transparent sequence audit trails to ensure responsible use.

To prevent unintended use, we claim that the code is not intended for clinical or therapeutic deployment. Additionally, users are encouraged to conduct proper validation before applying the code to any real-world biological or medical contexts.

